# SARS coronavirus vaccines protect against different coronaviruses

**DOI:** 10.1101/2021.06.01.446491

**Authors:** Tanushree Dangi, Nicole Palacio, Sarah Sanchez, Jacob Class, Lavanya Visvabharathy, Thomas Ciucci, Igor Koralnik, Justin Richner, Pablo Penaloza-MacMaster

**Affiliations:** Department of Microbiology-Immunology, Feinberg School of Medicine, Northwestern University, Chicago, IL 60611, USA; Department of Microbiology & Immunology, University of Illinois in Chicago, Chicago, IL 60612, USA; Division of Neuro-Infectious Diseases & Global Neurology; David H. Smith Center for Vaccine Biology and Immunology, University of Rochester, Rochester, NY 14642, USA; Department of Microbiology and Immunology, Center for Vaccine Biology and Immunology, University of Rochester, Rochester, NY 14642, USA.

**Author notes:** Correspondence: Justin Richner & Pablo Penaloza-MacMaster. These authors contributed equally (Tanushree Dangi, Nicole Palacio).

## Abstract

Although SARS-CoV-2 vaccines have shown efficacy against SARS-CoV-2, it is unclear if they can also protect against other coronaviruses that may infect humans in the future. Here, we show that SARS-CoV-2 vaccination in humans elicits cross-reactive antibodies against other coronaviruses. Our studies in mice demonstrate that SARS-CoV-2 vaccination protects against a common cold coronavirus, and that SARS-CoV-1 vaccination protects against SARS-CoV-2. Similarly, infection with a common cold coronavirus also conferred enhanced protection from subsequent infections with other coronaviruses. Mechanistically, both T cells and antibodies mediated cross-protection. This is the first direct demonstration that coronavirus-specific immunity can confer heterologous protection *in vivo*, providing a rationale for universal coronavirus vaccines.

**Highlights:**

1. SARS-CoV-2 vaccination elicits cross-reactive antibody against other coronaviruses in humans.
2. COVID-19 patients generate cross-reactive antibody against other coronaviruses.
3. A SARS-CoV-1 vaccine protects against SARS-CoV-2.
4. Prior coronavirus infections improve immune protection following heterologous coronavirus challenges.

## Introduction

Coronaviruses have garnered attention for their potential to cause pandemics. In less than 20 years, there have been outbreaks from at least three betacoronaviruses: Severe Acute Respiratory Syndrome associated coronavirus (SARS-CoV-1), Middle Eastern Respiratory Syndrome virus (MERS), and recently, SARS-CoV-2. Multiple SARS-CoV-2 vaccines have shown efficacy at preventing COVID-19, but whether they confer protection to other coronaviruses is unknown. It also remains unclear if infection with endemic coronaviruses confers protection against other coronaviruses. To answer these questions, we evaluated cross-protective immunity following coronavirus vaccination and coronavirus infection.

### SARS-CoV-2 vaccines induce cross-reactive antibody responses against other coronaviruses in humans

We first measured antibody responses following vaccination of humans with SARS-CoV-2 vaccines (Pfizer/BioNTek, Moderna and J&J). Plasma samples from human volunteers were obtained before vaccination, and at several time points after vaccination. As expected, vaccination of humans with SARS-CoV-2 vaccines resulted in increase in SARS-CoV-2 spike-specific antibodies (Fig. 1A). Importantly, the SARS-CoV-2 vaccines also induced an increase in SARS-CoV-1 spike-specific antibody (Fig. 1B). We also quantified antibody responses against the spike protein of OC43, which is an endemic coronavirus that causes common colds in humans. All patients had high levels of pre-existing antibody titers against OC43, but SARS-CoV-2 vaccination increased antibody titers against this endemic coronavirus (Fig. 1C). We then segregated our data, separating people who were infected with SARS-CoV-2 prior to vaccination from those who were not previously infected. Consistent with prior reports^1, 2^, a SARS-CoV-2 vaccine prime induced a substantial increase in SARS-CoV-2-specific antibody in people who were previously exposed to SARS-CoV-2 (Fig. 1D). A similar pattern was observed for SARS-CoV-1-specific antibody (Fig. 1E). Antibody responses to the more distant OC43 coronavirus were not increased further by the booster vaccine in unexposed individuals, and were higher in people who were previously infected (Fig. 1F). Taken together, these data showed that SARS-CoV-2 vaccination elicited cross-reactive antibody responses that were specific to other coronaviruses besides SARS-CoV-2.

**Figure 1.**
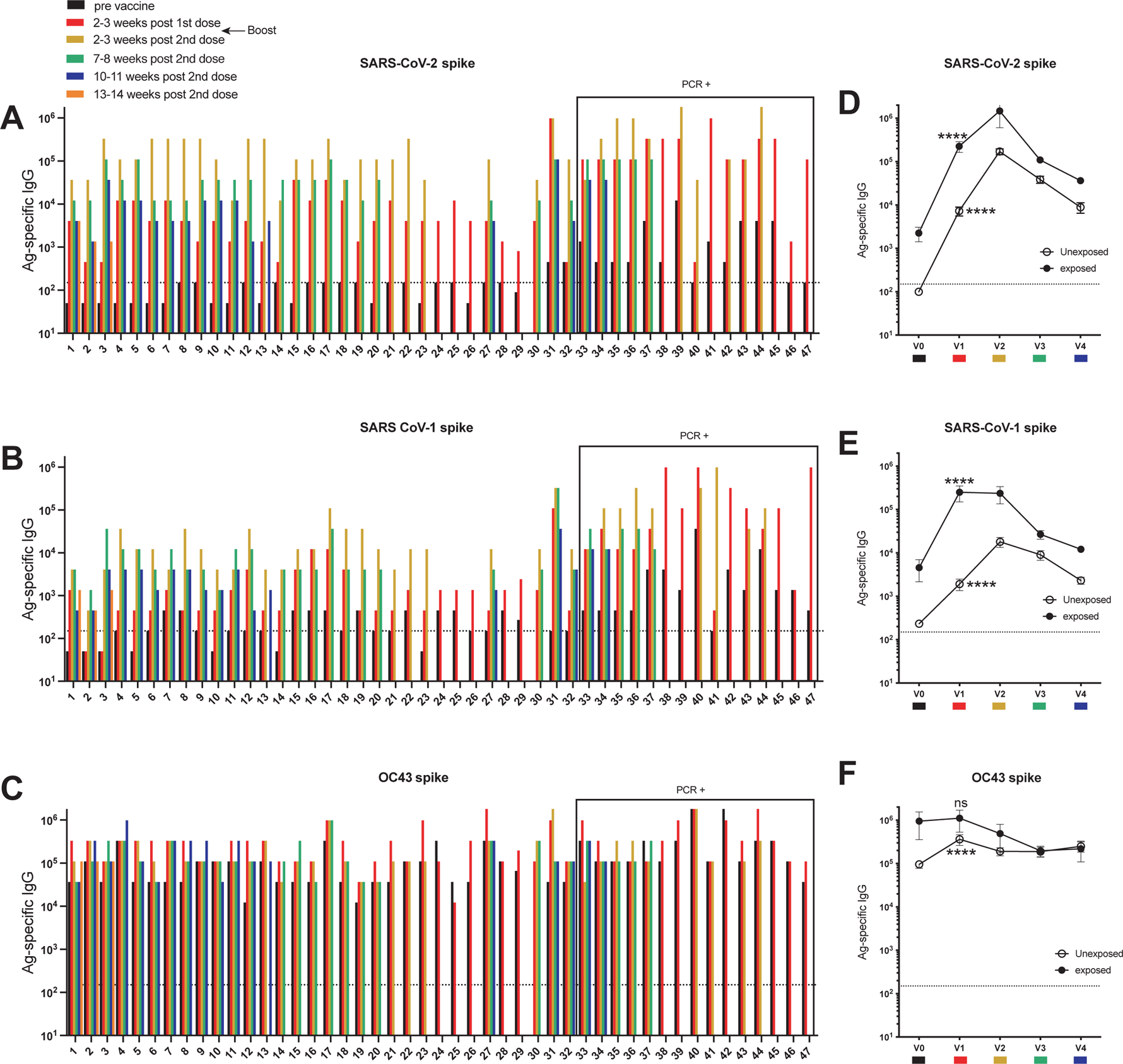
Cross-reactive antibody responses following SARS-CoV-2 vaccination. (A-D) Antibody responses after SARS-CoV-2 vaccination in humans. Participants 15-20, 25, 28 and 39 received the Moderna vaccine, participant 29 received the Johnson & Johnson (J&J) vaccine. The rest of participants received the Pfizer/BioNTek vaccine. Participants 27-30 are under different immunosuppressant regimens and did not stop their medication intake at the time of vaccination. Participants labeled as PCR+ tested positive for SARS-CoV-2 infection prior to vaccination. (**A**) SARS-CoV-2 spike-specific antibody responses. Samples 29 and 30 showed higher baseline SARS-CoV-2 spike-specific response. (**B**) SARS-CoV-1 spike-specific antibody responses. (**C**) OC43 spike-specific antibody responses. (D-F) Summary of antibody responses following vaccination in SARS-CoV-2 unexposed (open circles), and SARS-CoV- 2 exposed (closed circles) individuals. (**D**) Summary of SARS-CoV-2 spike-specific antibody responses. (**E**) Summary of SARS-CoV-1 spike-specific antibody responses. (**F**) Summary of OC43 spike-specific antibody responses. Antibody responses were evaluated by ELISA. Dashed lines represent limit of detection. In panels D-F, the numbers indicate matched P values comparing v0 (pre-vaccination) and v1 (week 2-3 post-prime) from each group by Wilcoxon test. ****, P <0.0001, ns, P > 0.05. Error bars represent SEM.

### COVID-19 patients show cross-reactive antibody responses against other coronaviruses

We then interrogated if cross-reactive antibodies could also be observed in COVID-19 patients. We compared antibody responses in plasma from PCR- confirmed COVID-19 patients admitted to the Northwestern Memorial Hospital, as well as healthy control plasma harvested before 2019. COVID-19 patients showed higher levels of SARS-CoV-2 spike-specific antibodies (Fig. 2A), as well as SARS-CoV-1 spike-specific (Fig. 2B) and OC43-specific (Fig. 2C) antibodies, relative to control individuals. These data demonstrate that COVID-19 patients develop cross-reactive antibody responses that recognize other coronaviruses.

**Figure 2.**
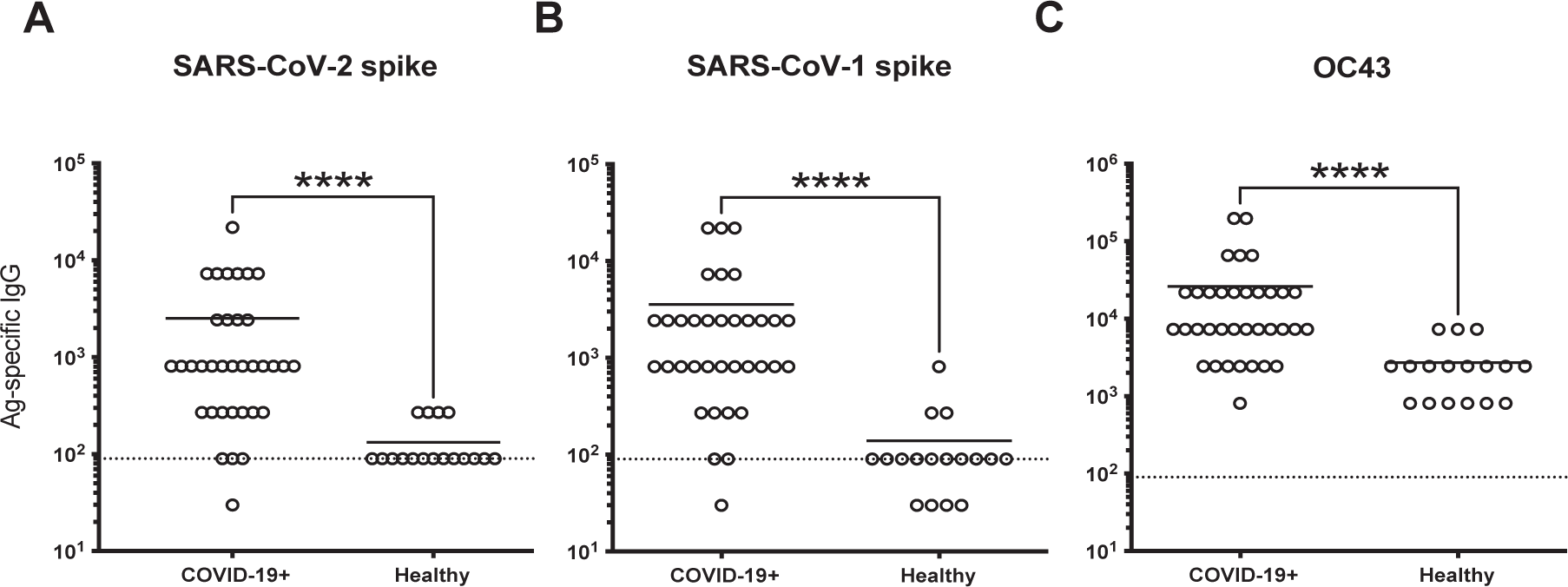
Cross-reactive antibody responses following SARS-CoV-2 infection in humans. Antibody responses after SARS-CoV-2 infection in symptomatic PCR-confirmed patients admitted to the Northwestern University Hospital. (**A**) SARS-CoV-2 spike-specific antibody responses. (**B**) SARS-CoV-1 spike-specific antibody responses. (**C**) OC43-specific antibody responses. OC43 infected cell lysates were used as coating antigen. Antibody responses were evaluated by ELISA. Dashed lines represent limit of detection. ****, P <0.0001 by Mann Whitney U Test.

### Characterization of cross-reactive antibody responses with multiple SARS-CoV-2 vaccine modalities

Our experiments above showed that SARS-CoV-2 vaccines induce antibody responses against heterologous coronaviruses in humans. We then interrogated whether this effect was generalizable to other vaccine platforms that have been used in humans. We primed C57BL/6 mice intramuscularly with various SARS-CoV-2 vaccines, including adenovirus-based, vesicular stomatitis virus (VSV) based, mRNA-based, RBD protein-based, spike protein-based, and inactivated virus-based vaccines. We boosted mice at approximately 3 weeks, and evaluated antibody responses at 2 weeks post-boost.

Consistent with our data in humans, vaccination of mice with an adenovirus vector expressing SARS-CoV-2 spike (Ad5-SARS-2 spike) resulted in potent antibody responses against SARS-CoV-2 and SARS-CoV-1, and a more modest but statistically significant increase in antibody responses against more distant coronaviruses, including OC43 and mouse hepatitis virus (MHV-1) (Fig. 3A). Cross-reactive antibody responses were also elicited by VSV-based, mRNA-based, RBD protein-based, spike protein-based, and inactivated virus-based vaccines (Fig. 3B-3F). Altogether, these data showed that multiple SARS-CoV-2 vaccine platforms are able to elicit cross-reactive antibody responses.

**Figure 3.**
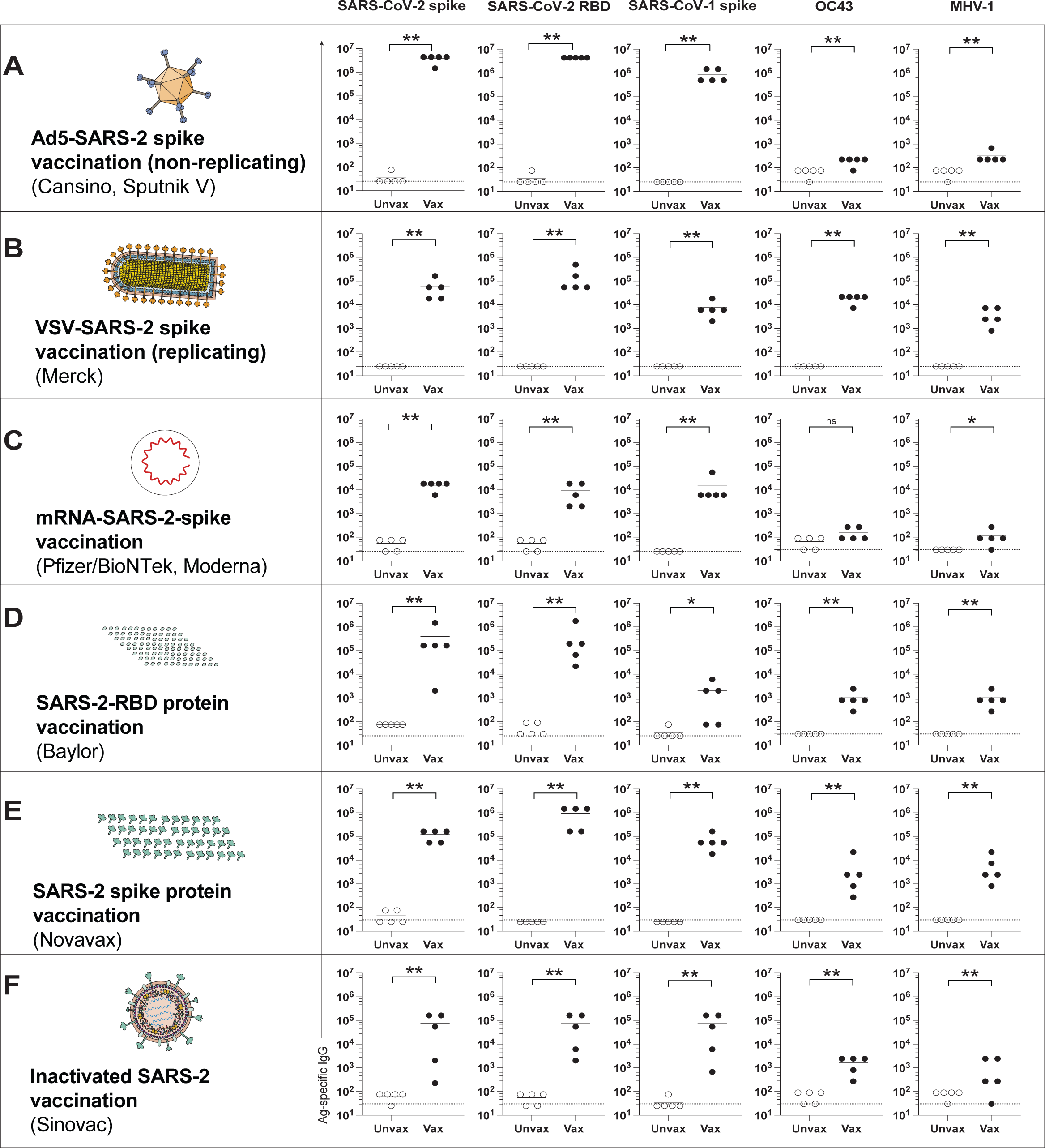
Cross-reactive antibody responses following SARS-CoV-2 vaccination in mice. (**A**) Antibody responses after Ad5-SARS-CoV-2 spike vaccination. (**B**) Antibody responses after VSV-SARS-CoV-2 spike vaccination. (**C**) Antibody responses after mRNA-SARS-CoV-2 spike vaccination. (**D**) Antibody responses after SARS-CoV-2 RBD vaccination. (**E**) Antibody responses after SARS-CoV-2 “whole” spike vaccination. (**F**) Antibody responses after inactivated SARS-CoV-2 vaccination. Mice were primed intramuscularly and boosted after 3 weeks (see Materials and Methods for vaccine dosing information). Antibody responses were evaluated by ELISA at week 2 post-boost. Experiments were done using wild type C57BL/6 mice, except for VSV-SARS- CoV-2 spike vaccination, which used k18-hACE2 (C57BL/6) mice. In each panel we indicate in parenthesis examples of clinically approved and experimental SARS-CoV-2 vaccines that are based on the same vaccine modality. Dashed lines represent limit of detection. Data are from 1 representative experiment with n=5/group; experiments were performed 2-3 times with similar results. *, P <0.05, **, P <0.01, ns, P > 0.05 by Mann Whitney U Test.

We then interrogated whether a vaccine against a different SARS coronavirus spike protein could also induce cross-reactive antibodies. Similarly, cross-reactive antibodies were also observed with an experimental SARS-CoV-1 spike vaccine developed in 2004, based on modified vaccinia Ankara (MVA-SARS-1 spike), which was previously shown to protect mice and macaques against a SARS-CoV-1 challenge^3, 4^ (Fig. 4A). Interestingly, sera from MVA-SARS-1- vaccinated mice partially neutralized SARS-CoV-2 pseudovirus *in vitro* (Fig. 4B-4D). These data show that immunization with a SARS-CoV-1 vaccine elicits cross-reactive neutralizing antibodies against SARS-CoV-2 and other coronaviruses.

**Figure 4.**
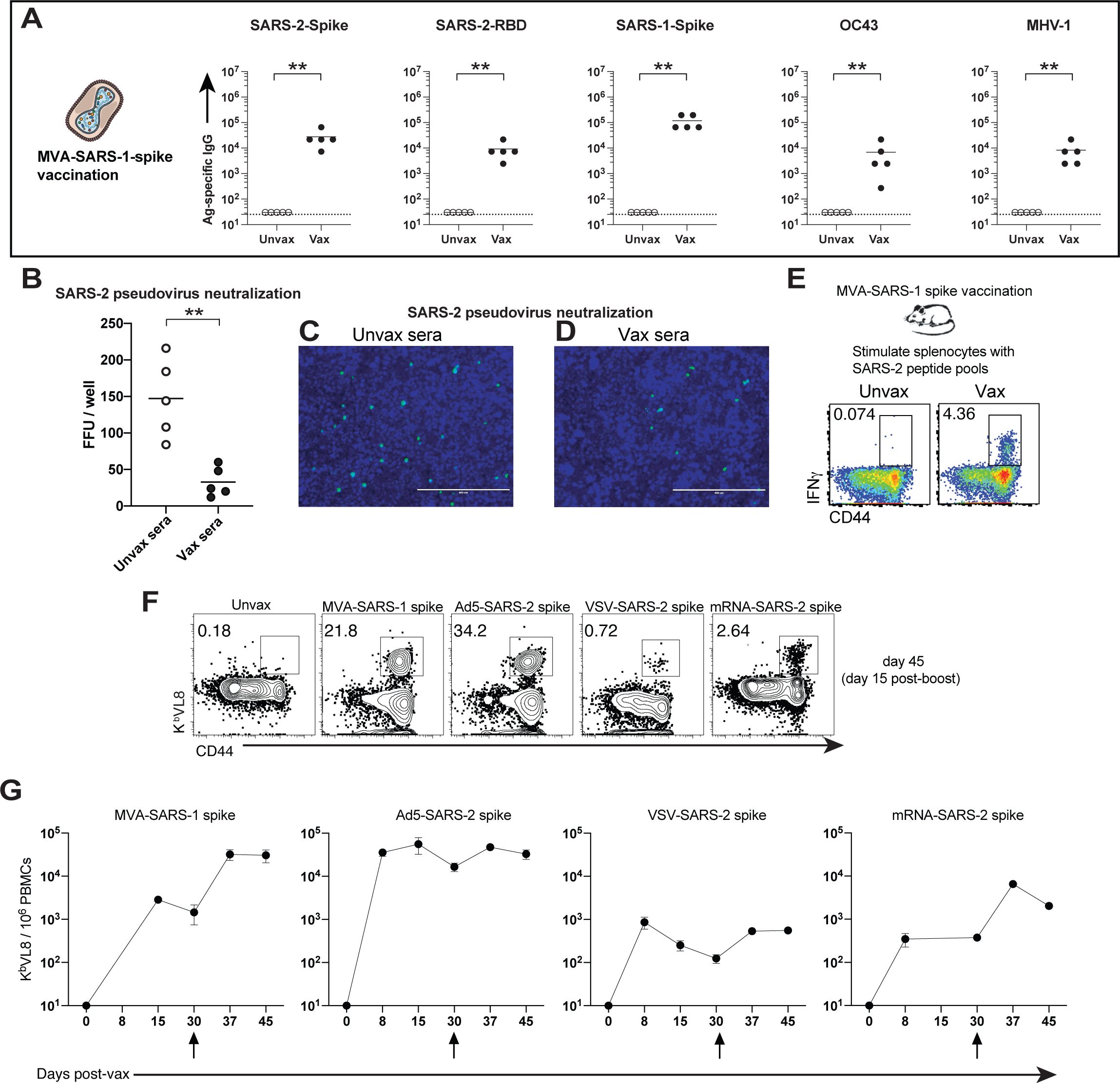
SARS-CoV-1 vaccination induces cross-reactive antibodies and T cells. (**A**) Antibody responses after MVA-SARS-CoV-1 spike vaccination. (**B**) SARS-CoV-2 pseudovirus neutralization assay. (**C**) Representative microscopy image of SARS-CoV-2 pseudovirus neutralization using sera from unvaccinated mice. (**D**) Representative microscopy image of SARS-CoV-2 pseudovirus neutralization using sera from SARS-CoV-1 vaccinated mice. (**E**) Representative FACS plots showing cross-reactive SARS-CoV-2 specific CD8 T cells in SARS-CoV-1 vaccinated mice. Cross-reactive CD8 T cells were detected by intracellular cytokine staining after 5 hr stimulation with SARS-CoV-2 spike overlapping peptide pools, in a 37°C 5% CO_2_ incubator. Cells are gated from total live CD8 T cells in spleen (week 2 post-boost). (**F**) Representative FACS plots showing cross-reactive (VNFNFNGL-specific) CD8 T cells in mice vaccinated with a SARS-CoV-1 vaccine, and various SARS-CoV-2 vaccines (Ad5-based, VSV-based and mRNA-based). Note that the K^b^ VNFNFNGL (K^b^ VL8) tetramer could be used to identify cross-reactive CD8 T cell responses among multiple vaccine platforms. Cells are gated from total live CD8 T cells in PBMCs (day 15 post-boost). (**G**) Summary of cross-reactive (VNFNFNGL-specific) CD8 T cells in mice vaccinated with a SARS-CoV-1 vaccine, and various SARS-CoV-2 vaccines. All mice were primed and boosted intramuscularly (see Materials and Methods for vaccine dosing information). Arrows in panel G indicate time of boosting. Experiments were done using wild type C57BL/6 mice, except for VSV-SARS-2 spike vaccination, which used k18- hACE2 (C57BL/6) mice. Dashed lines represent limit of detection. Data are from 1 representative experiment with n=5/group; experiments were performed 2-3 times with similar results. **, P <0.01 by Mann Whitney U Test.

Following a viral infection, viral control is facilitated by CD8 T cells. To measure cross-reactive CD8 T cell responses, we harvested splenocytes from mice that received the SARS-CoV-1 vaccine, and stimulated these cells with SARS-CoV-2 spike peptides (Table S1) for 5 hr, followed by intracellular cytokine stain (ICS) to detect cross-reactive (SARS-CoV-2 spike-specific) CD8 T cells. Interestingly, the SARS-CoV-1 vaccine elicited SARS-CoV-2 specific CD8 T cell responses (Fig. 4E), suggesting the presence of conserved CD8 T cell epitopes in SARS-CoV-1 and SARS-CoV-2. To identify cross-reactive CD8 T cell epitopes, we performed sequence alignment (fig. S1) followed by epitope mapping. We identified two highly conserved epitopes in the spike protein, in particular the VVLSFELL and VNFNFNGL epitopes, which are highly conserved among other SARS-like coronaviruses (fig. S1). These two epitopes were previously identified in a prior study in SARS-CoV-2 infected mice^5^. The VNFNFNGL CD8 T cell response has also been reported to be elicited after SARS-CoV-1 infection in C57BL/6 mice^6^, and we show that it is also immunodominant after SARS-CoV-2 vaccination (fig. S2A). Both VVLSFELL and VNFNFNGL were predicted to bind K^b^ by MHC-I epitope prediction algorithms (see Materials and Methods).

We reasoned that K^b^ VNFNFNGL tetramers could be used to track cross-reactive CD8 T cells following SARS-CoV-1 or SARS-CoV-2 vaccination. The protein vaccines and the inactivated vaccine did not generate a K^b^ VNFNFNGL (K^b^ VL8) CD8 T cell responses (not shown), likely because CD8 T cell priming is dependent on intracellular protein expression. However, K^b^ VL8 CD8 T cell responses were observed after vaccination with MVA-based, adenovirus-based, mRNA-based and VSV-based SARS-CoV-1 and SARS-CoV-2 vaccines (Fig. 4F). In particular, adenovirus-based and MVA-based vaccines seemed to generate the greatest K^b^ VL8 CD8 T cell response (Fig. 4F-4G).

We then performed single cell TCR-seq analyses to interrogate whether the conserved K^b^ VL8 response exhibited a biased TCR usage. We show that most of this conserved response contained a TCR composed of Vα7/Vβ11 (fig. S2B-S2D). We are currently using this single cell TCR sequencing information to develop a TCR transgenic mouse that could be used to study cross-reactive CD8 T cells among different sarbecovirus infections. Altogether, our data showed that a SARS-CoV-1 vaccine also generates antibody and T cell responses that recognize other coronaviruses. These data suggested that a SARS-CoV-1 vaccine could protect against heterologous coronavirus challenges, including SARS-CoV-2.

### A SARS-CoV-1 vaccine protects against a SARS-CoV-2 challenge

There are concerns about emerging SARS-CoV-2 variants and whether they could escape vaccine-elicited protection^7^. There are also worries that SARS- CoV-1 may spill over again in the human population. Thus, a critical question is whether SARS-CoV-2 vaccines could also protect against SARS-CoV-1, as well as other coronaviruses. To answer this simple question, we performed challenge experiments to evaluate whether coronavirus vaccines could protect against different coronaviruses. SARS-CoV-1 is a select agent, so we were not able to challenge SARS-CoV-2 vaccinated animals with SARS-CoV-1 in our BL3 facilities. Instead, we evaluated whether a SARS-CoV-1 vaccine could protect against a SARS-CoV-2 challenge. We immunized mice with a SARS-CoV-1 spike vaccine (MVA-SARS-1 spike), and then challenged mice intranasally with SARS-CoV-2. At day 5 post-challenge, we harvested lungs and measured viral loads by PCR. Strikingly, the SARS-CoV-1 vaccine conferred a 282-fold protection following a SARS-CoV-2 challenge (Fig. 5A). Improved control of SARS-CoV-2 was also observed at an earlier time post-challenge (day 3) (Fig. 5A). These data demonstrate that a SARS vaccine with only a 76% matched spike sequence could still confer robust immune protection.

**Figure 5.**
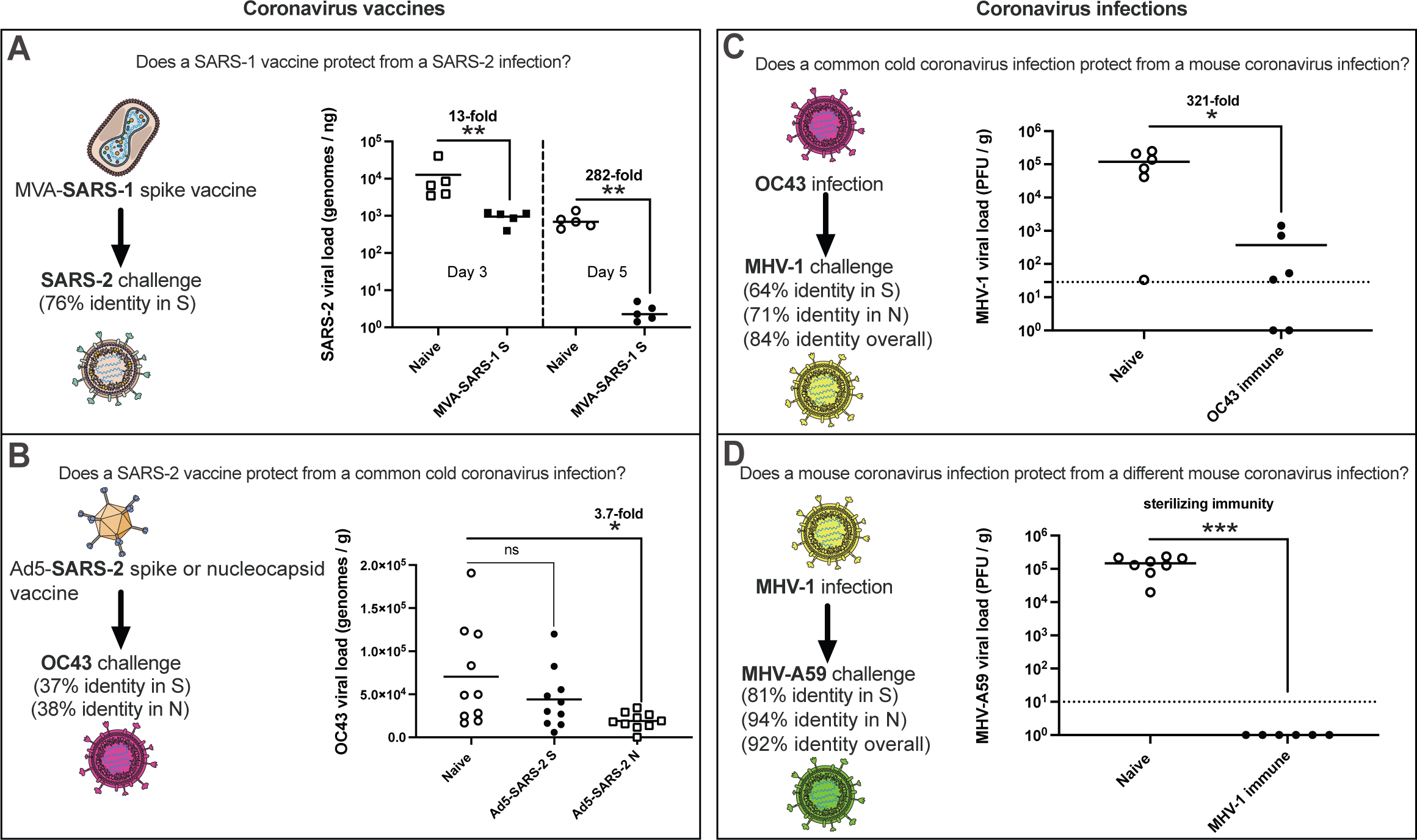
SARS-CoV-1 vaccination protects from SARS-CoV-2. (**A**) Viral loads after mouse-adapted SARS-CoV-2 MA10 challenge in MVA-SARS-CoV-1 vaccinated mice. Limit of detection is 0.007 genomes / ng. Data on the left (squares) are from one experiment in which mice were sacrificed at day 3 post-challenge. Data on the right (circles) are from one experiment in which mice were sacrificed at day 5 post-challenge. (**B**) Viral loads after OC43 challenge in Ad5- SARS-CoV-2 vaccinated mice (spike- versus nucleocapsid-based vaccine). Data are from day 5 post-challenge. Limit of detection is 27 genomes / g. (**C**) Viral loads after MHV-1 challenge in OC43-immune mice. Data are from day 5 post-challenge. (**D**) Viral loads after MHV-A59 challenge in MHV-1-immune mice. Data are from day 5 post-challenge. All mice were primed intramuscularly and boosted after 3 weeks (see Materials and Methods for vaccine dosing information). Mice were challenged intranasally after 2 weeks post-boost, and viral titers were measured in lungs using PCR (panels A-B) or plaque assays (panels C-D). We used BALB/c mice in panel A; C57BL/6 mice in panel B-C; and A/J mice in panel D. Dashed lines represent limit of detection. Data are from 2 experiments with n=3-5/group. Data from all experiments are shown. *, P <0.05, **, P <0.01, ***, P < 0.001, ns, P > 0.05 by Mann Whitney U Test.

Humans are constantly exposed to endemic coronaviruses. Our next question was whether a SARS-CoV-2 vaccine could confer protection against an endemic coronavirus (OC43), which shares less than 40% identity with SARS-CoV-2. We first immunized mice with a SARS-CoV-2 spike or nucleocapsid vaccine (both Ad5-based), and then challenged mice intranasally with OC43. At day 5 post-challenge, we harvested lungs and measured viral loads by PCR. The SARS-CoV-2 nucleocapsid vaccine (38% matched) conferred a 3.7-fold protection following this common cold coronavirus challenge (Fig. 5B). No significant heterologous protection was observed with the spike-based vaccine (37% matched) (Fig. 5B). Collectively, these data demonstrate that a SARS-CoV-2 coronavirus vaccine based on the nucleocapsid antigen can confer partial protection against a distant coronavirus challenge. In particular, the degree of heterologous protection appeared to be proportional to the genetic similarity between the vaccine antigen and the challenge antigen (Fig. 5A-5B).

### Prior coronavirus infections confer heterologous protection against different coronaviruses

Similar to our data with COVID-19 patients, coronavirus infections in mice also induced cross-reactive antibody responses. In particular, a common cold coronavirus infection elicited cross-reactive antibodies against SARS-CoV-2, SARS-CoV-1 and MHV-1 (fig. S3A). Cross-reactive antibody was also generated after an MHV-1 infection (fig S3B). We interrogated whether mice that had prior coronavirus infections were protected following heterologous coronavirus challenges. In our first model, we infected mice with the common cold coronavirus OC43, and after 4 weeks, we challenged these mice with MHV-1 (64% spike identity; 84% complete sequence identity). Importantly, OC43- immune mice exhibited a 321-fold protection against a heterologous MHV-1 challenge (Fig. 5C).

In our second model, we infected mice with MHV-1, and after 4 weeks, we challenged them with MHV-A59. Although these 2 coronaviruses have similar names, they are genetically distinct (81% spike identity; 92% complete sequence identity). Interestingly, MHV-1-immune mice exhibited sterilizing immune protection against a heterologous MHV-A59 challenge (Fig. 5D). These data demonstrate that coronavirus infections can confer partial or full protection against future infections with other coronaviruses. We also observed a pattern in which the degree of heterologous protection appeared to be proportional to the genetic similarity between the initial coronavirus and the subsequent coronavirus infection (Fig. 5C-5D).

### Mechanism: Both T cells and antibodies contribute to cross-protection

So far, we have shown that coronavirus vaccines and coronavirus infections induce cross-reactive immune responses in mice and humans. In addition, we showed that coronavirus vaccines and coronavirus infections confer protection against heterologous coronavirus challenges in mice. To analyze the specific contribution of T cells and antibody responses in cross-protection following heterologous coronavirus challenges, we performed challenge studies that allowed us to dissect the specific role of T cells and antibodies.

To assess the specific role of T cells, we immunized mice with a dendritic cell (DC)-based vaccine that induces SARS-CoV-2-specific T cell responses in the absence of antibody responses. This DC-based vaccine was prepared by coating DCs with short linear peptide pools derived from the SARS-CoV-2 spike, envelope, membrane and nucleocapsid proteins (DC-SARS-2 SEMN). After a week, mice were boosted with splenocytes coated with these same peptide pools, followed by a heterologous MHV-1 challenge to evaluate cross-protection (fig. S4A). This DC vaccine containing short linear peptides generated CD8 T cell responses (fig. S4B-S4C), but not antibody responses (fig. S4D). Interestingly, this SARS-CoV-2 T cell-based vaccine conferred robust protection against a heterologous MHV-A59 challenge (fig. S4E).

To evaluate the specific role of antibodies, we immunized mice with a SARS-CoV-1 spike vaccine, and then we harvested immune sera for adoptive transfer studies into naïve mice, followed by MHV-A59 challenge (fig. S5A). Although SARS-CoV-1 and MHV-A59 share only 30% identity in their spike proteins, SARS-CoV-1 spike-immune sera conferred partial protection in wild type recipient mice (fig. S5B) or *Ifnar1*-/- recipient mice (fig. S5C), which are highly susceptible to MHV-A59. In summary, our data showed that both T cell and antibody responses elicited by SARS coronavirus vaccination confer cross-protection against distant coronavirus challenges.

## DISCUSSION

SARS-CoV-2 has caused a major pandemic resulting in numerous deaths. Several SARS-CoV-2 vaccines have been deployed for human use, but it is unknown if these vaccines could also protect against other viruses, including pandemic or endemic coronaviruses. In this study, we show that SARS-CoV-2 vaccination in humans elicits cross-reactive antibody against SARS-CoV-1 and common cold coronavirus OC43. Our subsequent studies in mice demonstrate that a SARS-CoV-1 vaccine protects against a SARS-CoV-2 challenge, and that prior coronavirus infections can protect against subsequent infections with other coronaviruses. In our challenge models, both T cells and antibodies were able to mediate cross-protection.

Coronavirus vaccines has been previously shown to elicit cross-reactive antibodies^8–10^, but our study is the first to show *in vivo* heterologous protection by coronavirus vaccines. Our study suggests that cross-protection is proportional to the level of genetic conservation. For example, vaccination with a SARS-CoV-1 spike vaccine confers robust protection against a SARS-CoV-2 challenge (76% antigen-matched). On the other hand, vaccination with a SARS-CoV-2 spike vaccine did not confer significant protection against an OC43 challenge (only 37% antigen-matched). Similarly, a positive correlation between genetic similarity and heterologous protection were observed in the context of heterologous coronavirus infections. Therefore, our results partly mitigate concerns that current SARS-CoV-2 vaccines could become ineffective against SARS-CoV-2 variants, which are still 99% identical to the original SARS-CoV-2.

Moreover, our data suggest that infection may confer more robust cross-protective immunity than vaccination (Fig. 5). This could be explained by the high number of conserved antigens that are presented during a natural viral infection. However, most coronavirus vaccines are based only on the spike antigen, which is the least conserved protein of coronaviruses. It is possible that incorporation of other viral antigens may be necessary to develop universal vaccines with broad immune coverage.

Immune cross-reactivity in the context of coronavirus vaccination or coronavirus infection is based on genetic conservation of antigen sequences. A recent study showed that the SARS-CoV-2 nucleocapsid protein contains highly conserved amino acid sequences^11^. In particular, the PRWYFYYLGTGPEAGLPYG sequence is nearly identical in bat coronaviruses, human coronaviruses (e.g. OC43), and murine coronaviruses (e.g. MHV-A59 and MHV-1)^11^. Long linear stretches of conserved amino acid sequences may facilitate the priming of cross-reactive T cell responses. Additionally, the nucleocapsid is the most abundant structural protein in the coronavirus lifecycle, which may facilitate early detection by T cells ^12–14^. These peculiar features of the nucleocapsid protein may render it an optimal target for universal coronavirus vaccines, and may explain why in our studies a SARS-CoV-2 nucleocapsid-based vaccine, but not a spike-based vaccine, conferred partial protection against an OC43 challenge. Interestingly, vaccination with “whole” spike protein induced higher levels of cross-reactive antibody, relative to vaccination with RBD protein (Fig. 3D-3E). This is likely due to the high number of conserved epitopes in the “whole” spike protein.

Prior studies have shown that recent endemic coronavirus infections in humans are associated with less severe COVID-19^15^. In particular, young individuals show higher pre-existing immunity against seasonal coronaviruses, and this has been proposed to explain their improved clinical outcomes following SARS-CoV- 2 infection^16–20^. However, other studies have shown contradicting results^8^. Such discrepancy can be explained by the heterogeneity of immune histories and pre-existing conditions in humans, which can influence COVID-19 susceptibility, and the fact that those human studies were retrospective. Our study brings more clarity to this issue of heterologous protection mediated by prior coronavirus infections, as we utilize well-controlled mouse models with known immune histories. A limitation of our study is that we only evaluated heterologous immune protection at an early time post-vaccination or post-infection, and it is possible that cross-protective immunity declines over time. Future studies will determine whether cross-reactive antibodies are produced by plasma cells, or short-lived plasmablasts, which tend to be highly mutated and cross-reactive^21, 22^. In summary, we show that coronavirus vaccines and coronavirus infections can confer protection against other coronaviruses. These findings provide a framework for the rational design of universal coronavirus vaccines, and present the first definitive demonstration of heterologous immune protection following coronavirus vaccination or infection.

## AUTHOR CONTRIBUTIONS

P.P.M., J.R., T.D., and N.P designed and conducted the experiments. T.C. performed the scTCR-seq analyses. P.P.M., T.D., and N.P wrote the paper, with feedback from all other authors.

## Materials and Methods

### Human Subjects

All protocols used for subject recruitment, enrollment, blood collection, sample processing and immunological assays with human samples were approved by the Northwestern University Institutional review board (STU00212583). All participants voluntarily enrolled in the study by signing an informed consent form after receiving detailed information of the clinical study. Inclusion criteria consisted of being 18 years of age or greater, a SARS-CoV-2 infection and/or a scheduled vaccination for COVID-19, and ability and willingness to provide informed consent. Exclusion criteria consisted of less than 18 years of age or unwillingness or inability to provide informed consent. Enrollment started on August 2020 and is expected to be completed by August 2022. Subject population consisted of patients admitted to Northwestern Memorial Hospital and of residents across the Chicago area. Adults of different ages, races and ethnicities were included in the study. Subjects were de-identified by assigning a 4-letter study code, which will be used for the duration of the study. Hospital admitted patients were screened for SARS-CoV-2 infection by polymerase chain reaction (PCR) assay. Participants considered as exposed before vaccination had a positive PCR test for SARS-CoV-2 any time prior to vaccination. Blood was collected by phlebotomy using BD Vacutainer 10mL tubes containing sodium heparin. Anticoagulated blood was added to LeucoSep tubes (Greiner Bio) and plasma was separated by density gradient centrifugation. To protect subject’s identity, all samples were labeled with their assigned 4-letter study code and stored in principal investigator’s laboratory freezers.

### Mice, vaccinations, infections and challenges

6-8-week-old C57BL/6, BALB/c, A/J, *Ifnar*1-/- mice (on a C57BL/6 background) were used. For VSV-SARS-2 spike vaccinations, k18-hACE2 mice were used. Mice were purchased from Jackson laboratories (approximately half males and half females) and housed at the Northwestern University Center for Comparative Medicine (CCM) or the University of Illinois at Chicago (UIC). Mice were immunized intramuscularly (50 µl per quadriceps) with: adenovirus serotype 5 expressing SARS-CoV-2-spike protein (Ad5-SARS-2 spike; 10^9^ PFU), vesicular stomatitis virus expressing SARS-CoV-2-spike protein (VSV-SARS-2-spike; 10^7^ PFU), mRNA-based vaccine encoding SARS-CoV-2-spike protein (mRNA-SARS-2 spike; 5 µg), SARS-CoV-2 “whole spike” protein (SARS-2 spike; 100 µg), SARS-CoV-2 RBD protein (SARS-2 RBD protein; 100 µg), gamma-irradiated SARS-CoV-2 (inactivated SARS-2; 2.5x10^5^ PFU), and modified vaccinia Ankara expressing SARS-CoV-1 spike protein (MVA-SARS-1 spike; 10^7^ PFU).

We obtained Ad5-SARS-2 spike from the University of Iowa viral vector core (VVC-U-7643); VSV-SARS-2 spike from Dr. Sean Whelan (Washington University in St. Louis); and MVA-SARS-1 spike from the NIH Biodefense and Emerging Infections Research Resources Repository, NIAID, NIH (NR-623, originally developed by Dr. Bernard Moss^3^).

We synthesized mRNA vaccines encoding for the codon-optimized SARS-CoV-2 Spike protein from strain USA-WA1/2020. Constructs were purchased from Integrated DNA Technologies (IDT) and contained a T7 promoter site for in vitro transcription of mRNA, 5’ UTR and 3’ UTRs. The sequence of the 5’ and 3’ UTRs were identical to previous publications with a Dengue virus mRNA vaccine^23^. mRNA was synthesized from linearized DNA with T7 in vitro transcription kits from CellScript and following manufacturer’s protocol. RNA was generated with pseudouridine in place of uridine with the Incognito mRNA synthesis kit (Cat# C- ICTY110510). 5’ cap-1 structure and 3’ poly-A tail were enzymatically added. mRNA was encapsulated into lipid nanoparticles using the PNI Nanosystems NanoAssemblr Benchtop system. mRNA was dissolved in PNI Formulation Buffer (Cat# NWW0043) and was run through a laminar flow cartridge with GenVoy ILM (Cat# NWW0041) encapsulation lipids at a flow ratio of 3:1 (RNA in PNI Buffer : Genvoy ILM) at total flow rate of 12 mL/min to produce mRNA-LNPs. These mRNA-LNPs were characterized for encapsulation efficiency and mRNA concentration via RiboGreen Assay using Invitrogen’s Quant-iT Ribogreen RNA Assay Kit (Cat# R11490).

SARS-CoV-2 spike and RBD proteins used for vaccinations were produced at the Northwestern Recombinant Protein Production Core by Dr. Sergii Pshenychnyi using plasmids that were produced under HHSN272201400008C and obtained through BEI Resources, NIAID, NIH: Vector pCAGGS containing the SARS-related coronavirus 2, Wuhan-Hu-1 spike glycoprotein gene (soluble, stabilized), NR-52394 and receptor binding domain (RBD), NR-52309. Protein vaccines were administered with 1:10 AdjuPhos (Invivogen).

Inactivated SARS-CoV-2 was obtained from BEI resources, NIAID (SARS-related Coronavirus 2, isolate USA-WA1/2020, gamma-irradiated, NR-52287). MHV-1 was purchased from ATCC (VR-261) and OC43 was received from BEI (NR- 52725). MHV-A59 was a kind gift from Dr. Susan Weiss (University of Pennsylvania, Philadelphia, PA).

OC43 and MHV challenges: Mice were infected intranasally (25 µl per nostril) with OC43 (2x10^6^ PFU) or mouse hepatitis virus (MHV-1/MHV-A59; 10^6^ PFU). All mouse experiments with BL2 agents were performed with approval from the Northwestern University Institutional Animal Care and Use Committee (IACUC).

SARS-CoV-2 challenges: Mouse adapted SARS-CoV-2 (MA10) was kindly provided by Dr. Ralph Baric^24^. SARS-CoV-2 (MA10) was propagated and tittered on Vero-E6 cells. BALB/c mice were anesthetized with isoflurane and challenged via intranasal inoculation with 8x10^3^ FFU of SARS-CoV-2 (MA10). Lungs were isolated from mice at 5 days post infection and homogenized in PBS. RNA was extracted from lung homogenate using a Zymo Research Quick-RNA 96 Kit (R1052). Viral genomes were quantified via qPCR with N1 primer/probe kit from IDT (Cat. # 10006713). SARS-CoV-2 infections were performed at the University of Illinois at Chicago (UIC) following BL3 guidelines with approval by the UIC Institutional Animal Care and Use Committee.

### Protein-specific ELISA (SARS-CoV-2 spike, RBD, nucleocapsid; SARS-CoV- 1 spike; OC43 spike)

Antigen-specific total antibody titers were measured by ELISA as described previously (Dangi et al., 2020; Palacio et al., 2020). Briefly, 96-well flat-bottom MaxiSorp plates (Thermo Scientific) were coated with 1 µg/ml of respective protein, for 48 hr at 4°C. Plates were washed three times with wash buffer (PBS + 0.05% Tween 20). Blocking was performed with blocking solution (200 µl of PBS + 0.05% Tween 20 + 2% bovine serum albumin), for 4 hr at room temperature. 6 µl of sera were added to 144 µl of blocking solution in the first column of the plate, 1:3 serial dilutions were performed until row 12 for each sample, and plates were incubated for 60 min at room temperature. Plates were washed three times with wash buffer followed by addition of secondary antibody conjugated to horseradish peroxidase, goat anti-mouse IgG (Southern Biotech) diluted in blocking solution (1:1000) at 100 µl/well were added and incubated for 60 min at room temperature. For the ELISAs with human sera samples, goat anti-human IgG (H + L) conjugated with horseradish peroxidase (Jackson ImmunoResearch) was used. After washing plates three times with wash buffer, 100 µl/well of Sure Blue substrate (SeraCare) was added for 1 min. Reaction was stopped using 100 µl/well of KPL TMB Stop Solution (SeraCare). Absorbance was measured at 450 nm using a Spectramax Plus 384 (Molecular Devices).

SARS-CoV-2 spike and RBD proteins used for ELISA were produced at the Northwestern Recombinant Protein Production Core by Dr. Sergii Pshenychnyi using plasmids that were produced under HHSN272201400008C and obtained from BEI Resources, NIAID, NIH: Vector pCAGGS containing the SARS-related coronavirus 2, Wuhan-Hu-1 spike glycoprotein gene (soluble, stabilized), NR- 52394 and receptor binding domain (RBD), NR-52309. SARS-CoV-2 nucleocapsid protein was obtained through BEI Resources, NIAID, NIH (NR- 53797). SARS-CoV-1 spike protein was obtained through BEI Resources, NIAID, NIH (NR-722). OC43-spike protein was purchased from Sino Biologicals (40607- V08B).

### Virus-specific ELISA (OC43; MHV-1; MHV-A59)

Virus-specific ELISAs were performed as described earlier^25, 26^. In brief, 96-well flat-bottom MaxiSorp plates (Thermo Scientific) were coated with 100 µl/well of the respective viral lysate (OC43, MHV-1 or MHV-A59 infected cell lysates) diluted 1:10 in PBS, for 48 hr at room temperature. Plates were washed three times with wash buffer (PBS + 0.5% Tween 20) followed by blocking with blocking solution (200 µl/well of PBS + 0.2% Tween 20 + 10% FCS) for 2 hr at room temperature. 5 µl of sera were added to 145 µl of blocking solution in the first column of the plate, 1:3 serial dilutions were performed until row 12 for each sample, followed by incubation at room temperature for 90 min. Plates were washed three times with wash buffer, followed by addition of 100 µl/well of a secondary antibody conjugated to horseradish peroxidase, goat anti-mouse IgG (Southern Biotech) diluted in blocking solution (1:5000). Plates were incubated for 90 min at room temperature. Goat anti-human IgG (H + L) conjugated with horseradish peroxidase (Jackson ImmunoResearch) was used when ELISA was performed with human samples. After washing plates three times with wash buffer, 100 µl/well of Sure Blue substrate (SeraCare) was added for 8 min. Reaction was stopped using 100 µl/well of KPL TMB Stop Solution (SeraCare). Absorbance was measured at 450 nm using a Spectramax Plus 384 (Molecular Devices).

### Virus propagation

OC43 was propagated in an 80-90% confluent monolayer of HCT-8 cells (ATCC® CCL-244™) in T175 flasks at a multiplicity of infection (MOI) of 0.01 diluted in 5 mL of RPMI supplemented with 2% FBS, 1% penicillin/streptomycin, and 1% L-glutamine. Infected cells were incubated at 33°C for 2 hr in a humidified 5% CO2 incubator. After incubation, flasks were supplemented with 20 ml of 2% RPMI and incubated for 5 days at 33°C in a CO_2_ incubator. MHV- A59 and MHV-1 were expanded in 17CL-1 cells following a prior protocol^27^.

### OC43 and MHV quantification by plaque assay

For MHV quantification, L2 cells (10^6^ cells per well) were seeded into 6-well plates in 10% DMEM (10% FBS, 1% penicillin/streptomycin, and L-glutamine). After 2 days, when cells reached ∼100% confluency, media were removed. 10- fold serial dilutions of viral stock or homogenized lung were prepared in 1% DMEM (1% FBS, 1% penicillin/streptomycin, and L-glutamine), added to wells, and incubated at 37°C for 1 hr, gently rocking plates every 10 minutes. After incubation, 3.5 mL of 1% agarose diluted 1:1 with 20% 2X-199 media (2X-199 media supplemented with 20% FBS, 1% penicillin/streptomycin, and L-glutamine) was overlaid onto the monolayer and the plates were incubated at 37°C 5% CO_2_ for 2 days. On day 2, the agar overlay was removed gently and the monolayer was stained with 1% crystal violet for 15 minutes. After staining, the crystal violet was aspirated, plates were washed once with 2 ml water per well, and then dried to visualize plaques. Quantification of OC43 stocks for challenge studies was similar to the quantification of MHV-A59^27^ except that 5 ml of agar overlay was added on an infected monolayer of L2 cells and incubated at 33°C in CO_2_ incubator for 5-6 days. Monolayer was stained with 1% crystal violet and plaques quantified by manual counting. For viral load quantification in lung, tissue was collected in round-bottom 14-ml tubes (Falcon) containing 2 ml of 1% FBS DMEM. Tissues were ruptured using a Tissue Ruptor homogenizer (Qiagen). Homogenized tissues were clarified using a 100-µm strainer (Scientific Inc.) to remove debris, and clarified tissue lysates were used for plaque assay.

### Quantification of OC43 by RT-PCR

Lungs were isolated from mice and homogenized in 1% FBS DMEM. RNA was extracted from lung homogenate using PureLink Viral RNA/DNA Mini kit (Invitrogen), according to the manufacturer’s instructions. OC43 viral loads in lungs were determined using one-step quantitative real-time reverse transcriptase polymerase chain reaction (qRT-PCR). RT-PCR was performed using OC43-nucleocapsid specific TaqMan primers and a probe 18 fluorescent-labelled with a 5’-FAM reporter dye and 3’-BHQ quencher (IDT) and AgPath-ID™ One-Step RT-PCR kit (AgPath AM1005, Applied Biosystems) on an ABI QuantStudio 3 platform (Thermo Fisher). Each sample was tested in duplicates in 25 µl reactions containing 12.5 µl of a 2X RT-PCR buffer, 1 µl of 25X RT-PCR enzyme mix provided with the AgPath kit, 0.5 µl (450 nM) forward primer, 0.5 µl (450 nM) reverse primer, 0.5 µl (100 nM) of probe, and 10 µl RNA. In parallel, each sample was also tested for beta-actin gene as an internal control to verify RNA extraction quality using mouse beta-actin-specific Taqman primers/probe labelled with 5’-FAM and 3’-BHQ (IDT). Thermal cycling involved reverse transcription at 45°C for 10 min, denaturation at 95°C for 15 min, followed by 45 cycles of amplification (15 sec at 95°C and 1 min at 60°C.) To avoid cross-contamination, single use aliquots were prepared for all reagents including primers, probes, buffers, and enzymes.

### Quantification of SARS-CoV-2 by RT-PCR

Lungs were isolated from mice and homogenized in PBS. RNA was extracted from lung homogenate using a Zymo Research Quick-RNA 96 Kit (R1052). Viral genomes were quantified via RT-qPCR with the TaqMan RNA-to-Ct 1-step kit (ThermoFisher, Cat # 4392653) and primer/probe sets with the following sequences: Forward 5’ GAC CCC AAA ATC AGC GAA AT 3’, Reverse 5’ TCT GGT TAC TGC CAG TTG AAT CTG 3’, Probe 5’ ACC CCG CAT TAC GTT TGG TGG ACC 3’ (Integrated DNA Technologies, Cat # 10006713). A SARS CoV-2 copy number control was obtained from BEI (NR-52358) and used to quantify SARS-CoV-2 genomes.

### Dendritic cell (DC) vaccination

We followed a protocol similar to that from a prior publication^28^, but using DC2.4 cells instead of BMDCs. In brief, after thawing DC2.4 cells were passaged 3 times in 10% RPMI (10% FBS, penicillin/streptomycin, and L-glutamine). When the monolayer reached ∼70% confluency, the DCs were stimulated with LPS (100 ng/ml; Sigma Aldrich) and incubated for 1 day in a 5% CO_2_ incubator at 37°C. The next day, a cocktail of overlapping peptide pools (SARS-CoV-2 spike, envelope, membrane, and nucleocapsid: SARS-2-SEMN) was added to the DCs for 2 hr (1 µM/peptide). DCs were washed 5x with PBS, and 10^6^ DCs were injected into C57BL/6 mice intravenously via the lateral vein. At day 7 post-prime, splenocytes from a naïve mouse were coated with the same peptide pools and incubated for 2 hr, followed by washing 5x with PBS. Mice were then boosted with 20x10^6^ peptide-coated splenocytes intraperitoneally. DMSO (vehicle) coated DCs and spleen cells were used as controls.

The SARS-CoV-2 <u>spike</u> overlapping peptide pools were obtained from BEI resources, NIH (NR-52402). The SARS-CoV-2 <u>envelope</u> overlapping peptide pools were obtained from BEI resources, NIH (NR-52405). The SARS-CoV-2 <u>membrane</u> overlapping peptide pools were obtained from BEI resources, NIH (NR-52403). The SARS-CoV-2 <u>nucleocapsid</u> overlapping peptide pools were obtained from BEI resources, NIH (NR-52404).

### Reagents, flow cytometry, and equipment

Dead cells were gated out using Live/Dead fixable dead cell stain (Invitrogen). The SARS-CoV-2 spike overlapping peptide pools obtained from BEI resources, NIH (NR-52402) were used for intracellular cytokine staining. Biotinylated MHC class I monomers (Kb VNFNFNGL, abbreviated as Kb VL8) were obtained from the NIH tetramer facility at Emory University. Cells were stained with fluorescently-labelled antibodies against CD44 (IM7 on Pacific Blue), CD8α (53- 6.7 on PerCP-Cy5.5), IFNγ (XMG1.2 on APC). Fluorescently-labelled antibodies were purchased from BD Pharmingen, except for anti-CD44 (which was from Biolegend). Flow cytometry samples were acquired with a Becton Dickinson Canto II or an LSRII and analyzed using FlowJo (Treestar).

### SARS-CoV-2 pseudovirus neutralization assays

A SARS-CoV-2 pseudovirus was generated by transfection of HEK-293T cells with a pCAGGS vector expressing the SARS-CoV-2 spike glycoprotein (BEI resources, NIAID, NIH: NR-52310). 24 hr later, transfected cells were infected with VSVΔG*G-GFP at a multiplicity of infection (MOI) of 0.5. After 24 hr, GFP foci were visualized, and the supernatant was harvested and passed through a 0.45 μM filter. This SARS-CoV-2 pseudovirus was concentrated using an Amicon Ultra-15 filter (UFC910024, Sigma-Aldrich), and then stored at −80°C. Titers were measured by infecting HEK-293T-hACE2 cells and counting GFP foci under a fluorescence microscope after 24 hr.

The SARS-CoV-2 pseudovirus neutralization assay was performed by mixing serial dilutions of MVA-SARS-CoV-1 immune mice sera (or naïve sera) with 200 FFU of SARS-CoV-2 pseudovirus in a 96-well plate and incubating for 2 hr. After incubation, 100 µl of the sera-virus mixture was transferred to a half area 96-well plate containing HEK-293T-hACE2 cells. The next day, GFP foci were counted in each well under a fluorescent microscope.

### MHC-I binding predictions

The MHC-I binding predictions were made on 5/17/2021 using the IEDB analysis resource NetMHCpan (ver. 4.1) tool^29^, at http://tools.iedb.org/mhci/.

### Single cell TCR-Seq Data Acquisition and Analysis

C57BL/6 mice were immunized intramuscularly with 10^9^ PFU of Ad5-SARS-2 spike, and at day 28, splenic CD8 T cells were MACS-sorted using negative selection (STEMCELL). Purified CD8 T cells were stained with K^b^ VL8, Live/Dead stain, and flow cytometry antibodies for CD8 and CD44. Live, CD8+, CD44+, K^b^ VL8+ cells were FACS-sorted to ∼99% purity on a FACS Aria cytometer (BD Biosciences) and delivered to the Northwestern University NU-Seq core for scTCR-seq using Chromium NextGem 5’ v2 kit (10X Genomics). Once the library was sequenced, the output file in BCL format was converted to fastq files and aligned to mouse genome in order to generate a matrix file using the Cell Ranger pipeline. These upstream QC steps were performed by Drs. Ching Man Wai and Matthew Schipma at the Northwestern University NUSeq core. TCR analyses were performed using the scRepertoire package^30^. Only cells expressing both TCRα and TCRβ chains were selected. For cells with more than 2 TCR chains, only the top 2 expressed chains were used.

### Statistical analyses

Most statistical analyses used the Mann-Whitney test, unless specified otherwise in the figure legend. Dashed lines in ELISA/plaque assay figures represent the limit of detection. Data were analyzed using Prism (Graphpad).

## Acknowledgments

We thank Dr. Thomas Gallagher for comments and suggestions. Our study was possible with a grant from the National Institute on Drug Abuse (NIDA, DP2DA051912) and a grant from the Emerging and Re-Emerging Pathogens Program (EREPP) to P.P.M.

## Supplemental Table Legends

Supplemental Table 1. SARS-CoV-2 spike overlapping peptide pools spanning 1273 amino acids. Each individual peptide consisted of 13-17 amino acids, with 10 amino acid overlaps.

## Supplemental Figure Legends

**Supplemental Figure 1.**
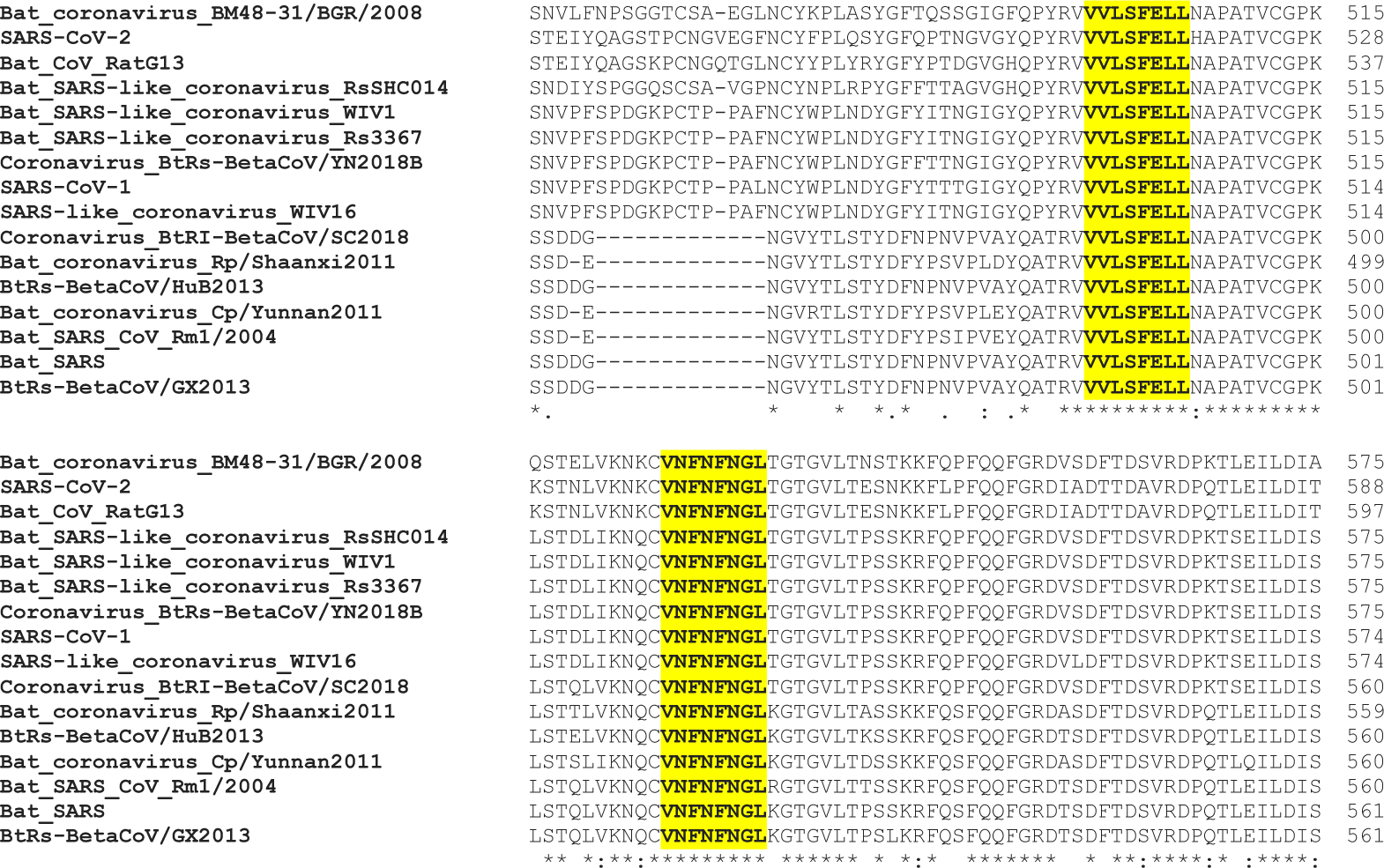
Gene alignment of conserved epitopes across various bat coronaviruses. Clustal Omega Multiple Sequence Alignment of SARS-CoV-1, SARS-CoV-2, and other betacoronaviruses, showing conservation of the VVLSFELL and VNFNFNGL epitopes.

**Supplemental Figure 2.**
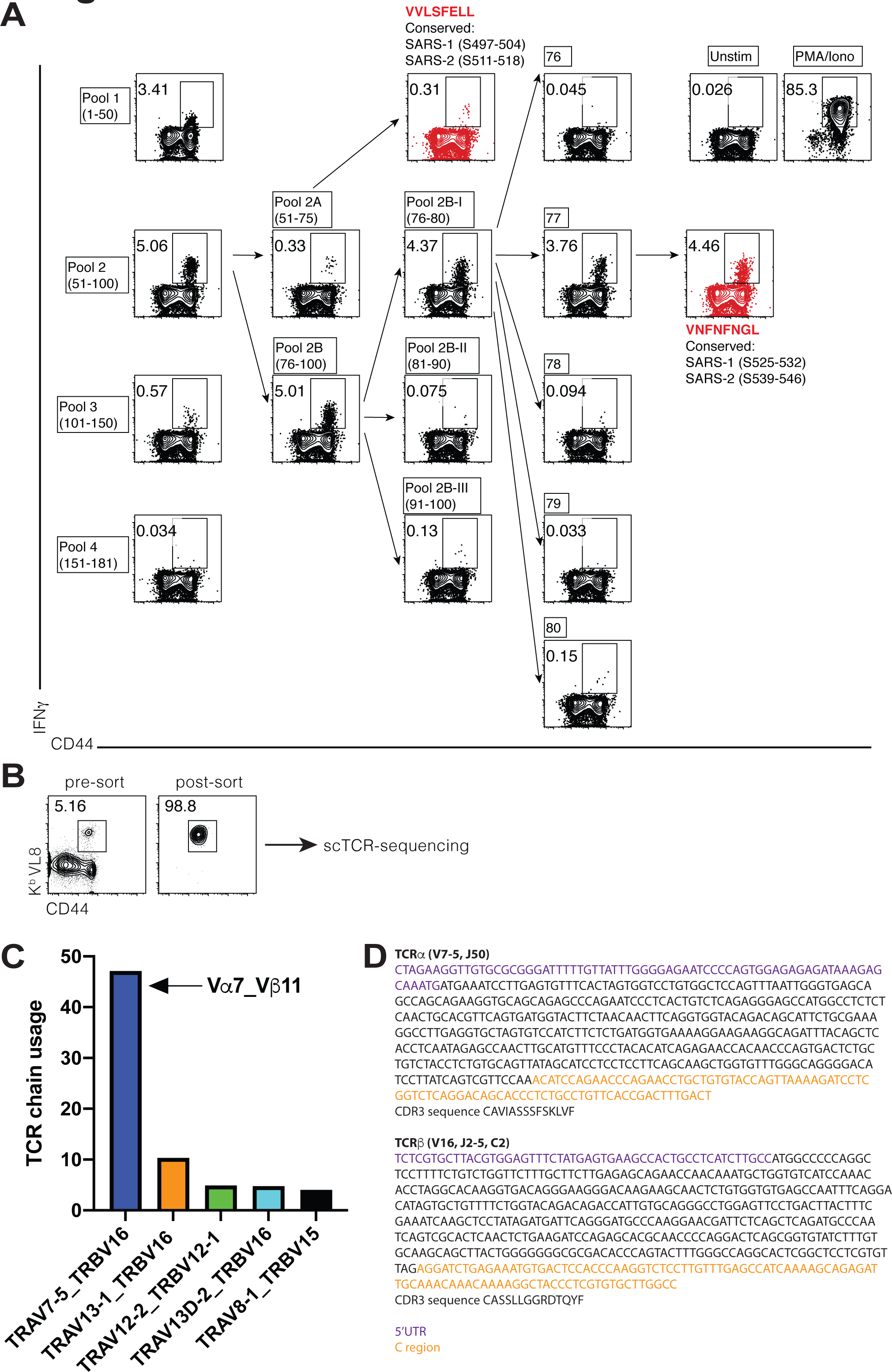
Mapping of conserved CD8 T cell epitopes following SARS-CoV-2 vaccination. (**A**) Epitope mapping using Ad5-SARS-2 spike immune splenocytes (week 2 post-boost), stimulated with overlapping SARS-CoV-2 spike peptide pools for 5 hr at 37°C in a CO_2_ incubator. This study identified 2 potential K^b^ binding epitopes that are highly conserved among multiple coronaviruses, a subdominant VVLSFELL epitope and a dominant VNFNFNGL epitope. (**B**) K^b^ VNFNFNGL (K^b^ VL8) tetramer+ CD8 T cells were FACS-sorted for TCR-sequencing (week 4 post-prime). (**C**) Top 5 TCR usages (percent of total VL8-specific). (**D**) TCR sequences in VL8-specific CD8 T cells. Data from panel A represent a pooled sample from 5 spleens. Data from panels B-D are from 1 experiment with 1 mouse.

**Supplemental Figure 3.**
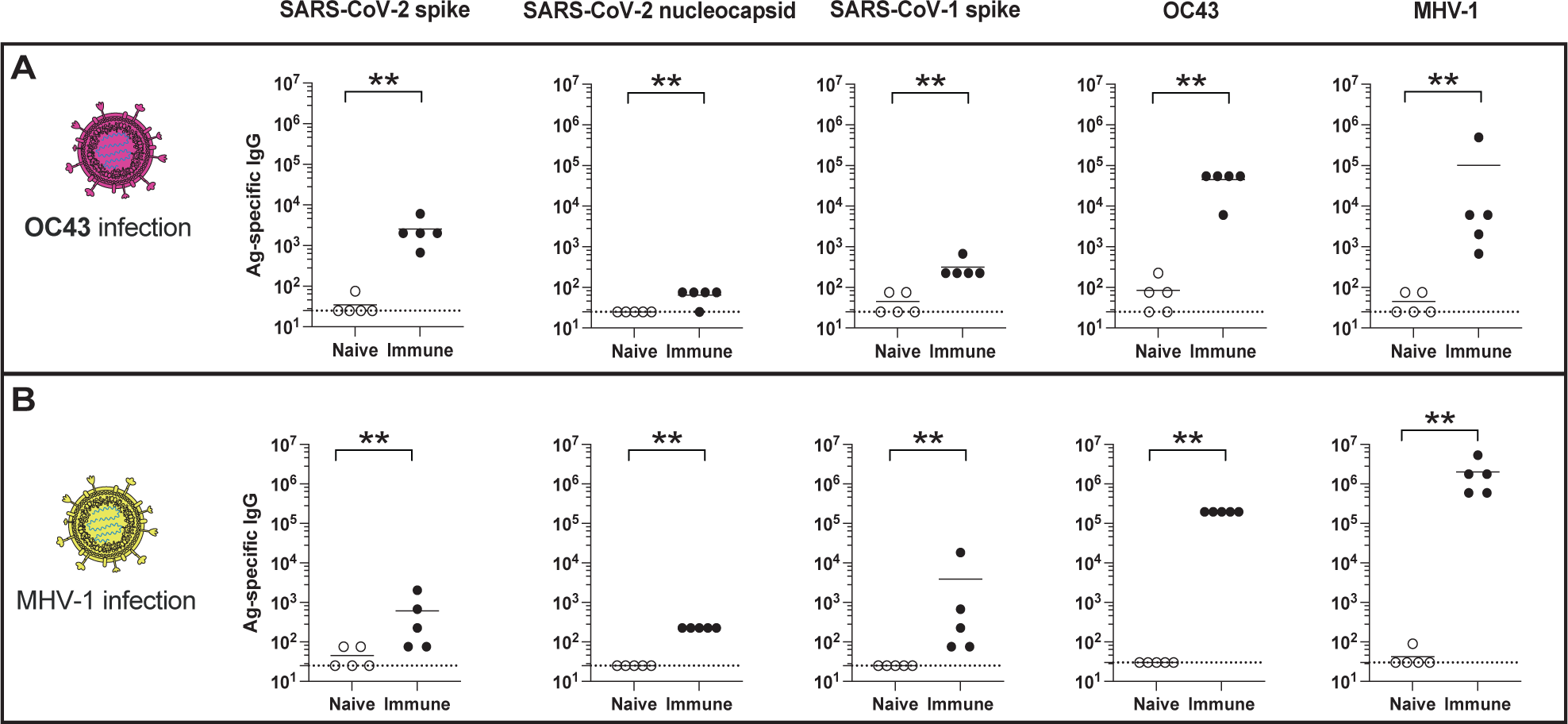
Cross-reactive antibody responses following coronavirus infections in mice. (**A**) Antibody responses after infection with OC43 common cold coronavirus. (**B**) Antibody responses after infection with MHV-1 coronavirus. Mice were infected intranasally and boosted after 3 weeks (see Materials and Methods for virus dose information). Antibody responses were evaluated by ELISA at week 2 post-boost. Experiments were done using wild type C57BL/6 mice. Dashed lines represent limit of detection. Data are from 1 representative experiment with n=5/group; experiments were performed 2 times with similar results. **, P <0.01 by Mann Whitney U Test.

**Supplemental Figure 4.**
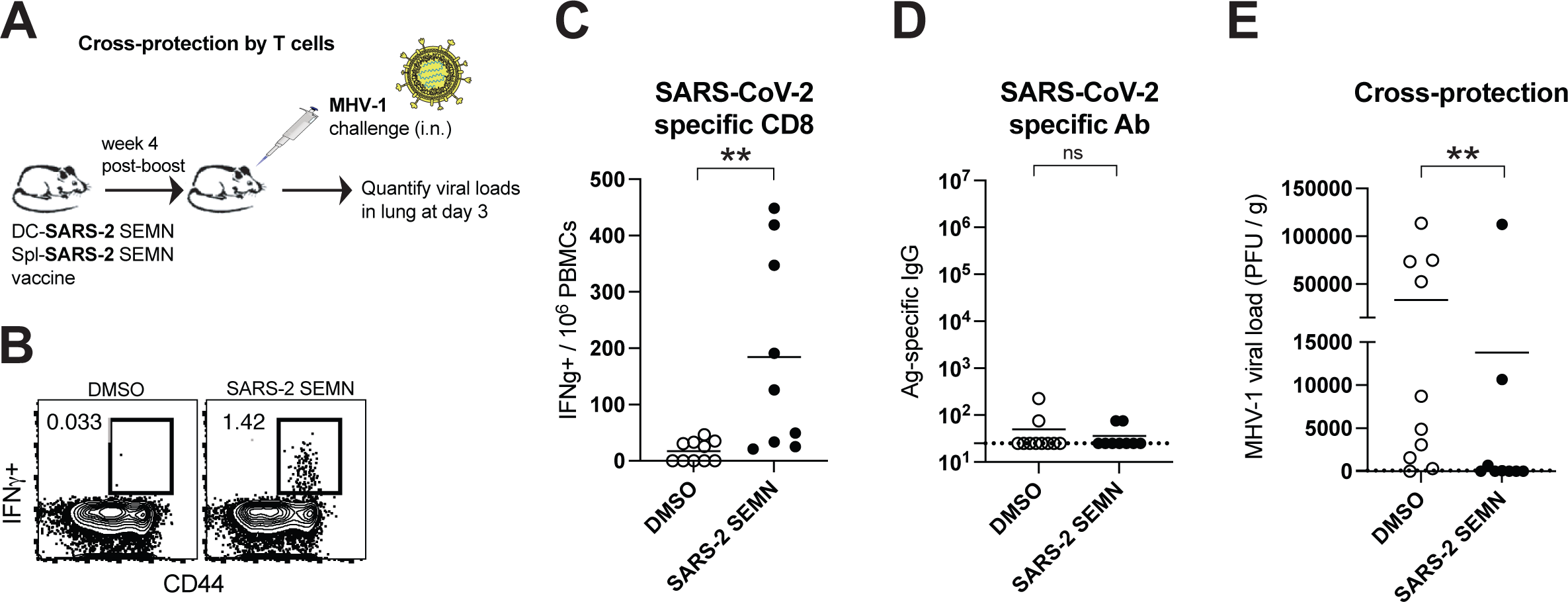
A T cell-based SARS vaccine protects against a distant coronavirus challenge. (**A**) Experiment outline. C57BL/6 mice received DC2.4 cells coated with SARS-CoV-2 peptide pools derived from <u>s</u>pike, <u>e</u>nvelope, <u>m</u>embrane, and <u>n</u>ucleocapsid proteins (DC-SEMN). After a week, mice were boosted with splenocytes coated with these same peptide pools. DMSO vehicle-coated cells were used as controls (see Materials and Methods). A week after boost, mice were challenged intranasally with MHV-1, and viral loads were assessed in lungs at day 3 by plaque assays. (**B**) Representative FACS plots showing SARS-CoV-2 SEMN-specific CD8 T cells. SARS-CoV-2 specific T cells were detected by intracellular cytokine staining after 5 hr stimulation with SARS- CoV-2 overlapping peptide pools (SEMN), in a 37°C 5% CO_2_ incubator. Cells are gated from total live CD8 T cells in spleen (week 1 post-boost). (**C**) Summary of SARS-CoV-2 specific CD8 T cell responses. (**D**) SARS-CoV-2 spike-specific antibody. (**E**) Viral loads after MHV-1 challenge. SARS-CoV-2 and MHV-1 share ∼38% identity in SEMN. Dashed lines represent limit of detection. Experiment was performed two times, with n=4-5 mice per group, per experiment. Data from all experiments are shown. P value is indicated (Mann Whitney U Test). **, P <0.01, ns, P > 0.05 by Mann Whitney U Test.

**Supplemental Figure 5.**
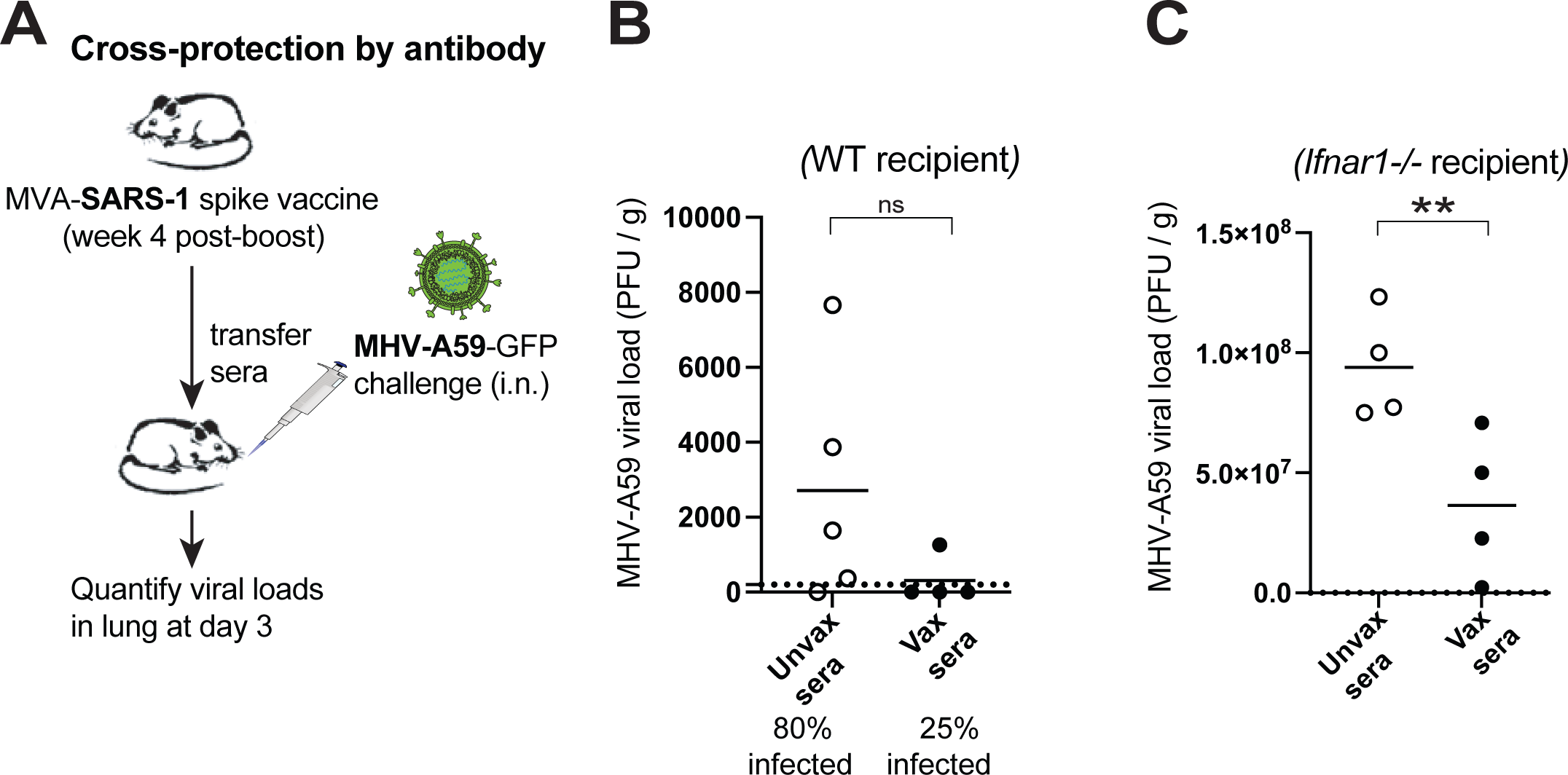
Antibody responses elicited by a SARS vaccine protect against a distant coronavirus challenge. (**A**) Experiment outline. C57BL/6 mice were vaccinated intramuscularly with MVA-SARS-CoV-1 spike and then boosted after 3 weeks. Mice were bled 2 weeks after boost to harvest immune sera, and 0.5 μL sera were transferred intraperitoneally to naïve recipient C57BL/6 mice. On the next day, mice were challenged intranasally with MHV-A59, and viral loads were assessed in lungs at day 3 by plaque assays. This passive immunization experiment was performed two times; the first experiment utilized wild type recipients and the second experiment utilized *Ifnar1*- /- recipients, which are highly susceptible to MHV-A59. (**B**) Protection against an MHV-A59 challenge in wild type mice. (**C**) Protection against an MHV-A59 challenge in *Ifnar1*-/- mice. Data from all experiments are shown. **, P <0.01, ns, P > 0.05 by Mann Whitney U Test.

